## Supplemental Figures and Table for "Structured connectivity in the output of the cerebellar cortex"

### Supplementary Material

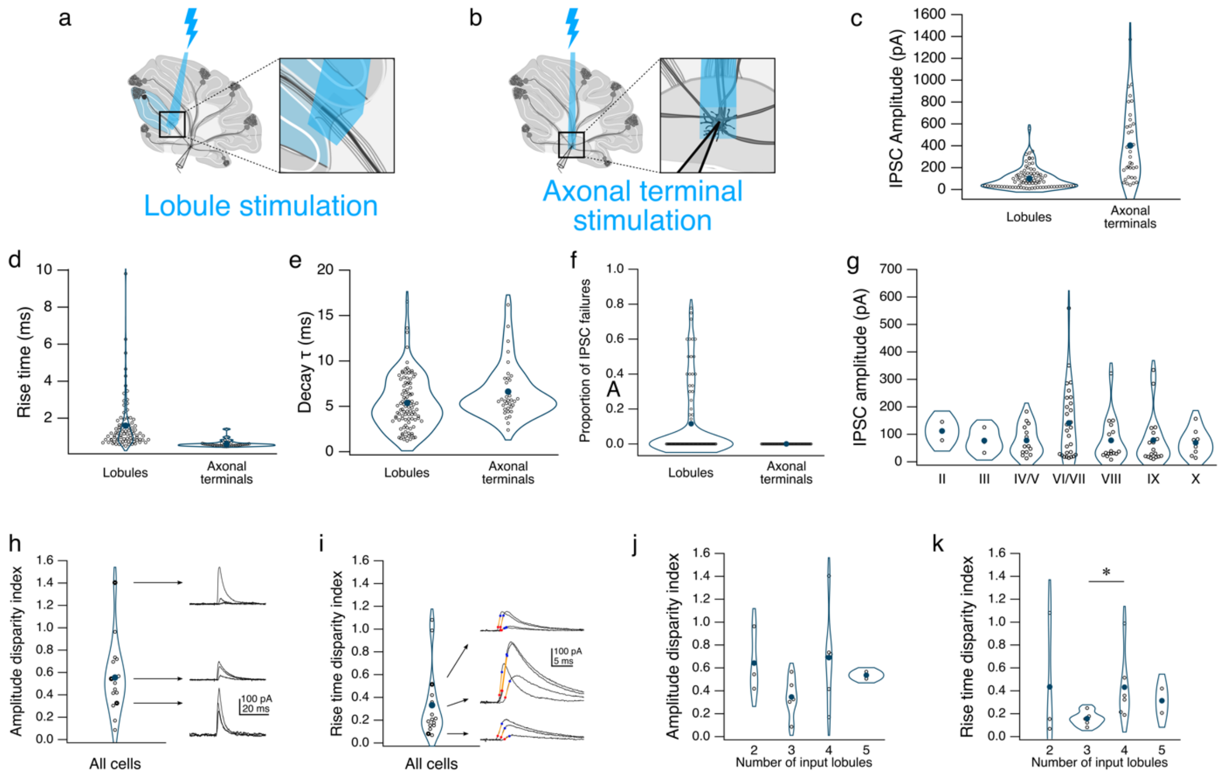


**Supplementary Figure 1**: Properties of inhibitory post-synaptic currents (IPSCs) following Purkinje cell stimulation. **a**, Cartoon showing area of photostimulation for “lobule stimulation,” and **b**, “axonal terminal stimulation.” **c-f**, IPSC amplitudes, rise times, weighted decay τ time constants, and failures evoked following “lobule” or “axonal terminal” stimulation. **g**, IPSC amplitudes by lobule. **h-i**, Average IPSC amplitude disparity index for all cells with input from > 1 lobule: 0.55 ± 0.08, and average rise time disparity index for all cells with input from > 1 lobule: 0.33 ± 0.08. Traces to the right of each graph displaying examples of IPSCs with different disparity index values. **j-k**, Disparity indices for IPSC amplitudes and rise times by number of lobule inputs. *$P < 0.05$. Unlabeled comparisons = not significantly different.

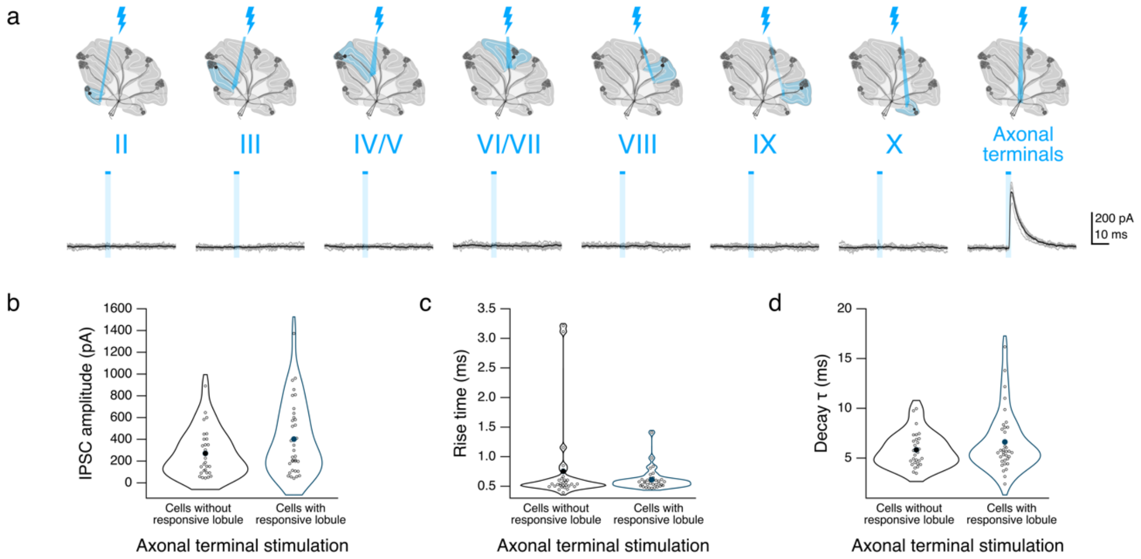


**Supplementary Figure 2**: Responses from cells with non-responsive lobules after lobule stimulation. **a**, Top, stimulation protocol as described in **Fig. 1b**. Bottom, example traces following lobule stimulation (left) and axonal terminal stimulation (right). **b**, Average IPSC amplitude for cells without responsive lobules versus cells with responsive lobules (Fig. 1) following axonal terminal stimulation. **c**, Rise times for cells without responsive lobules versus cells with responsive lobules following axonal terminal stimulation. **d**, Weighted decay τ time constants of IPSCs from cells without responsive lobules versus cells with responsive lobules. IPSC amplitudes, rise times, and decay τ time constants compared between cells with and without responsive lobules using non-parametric multiple comparison Mann-Whitney *U* test. Unlabeled comparisons = not significantly different.

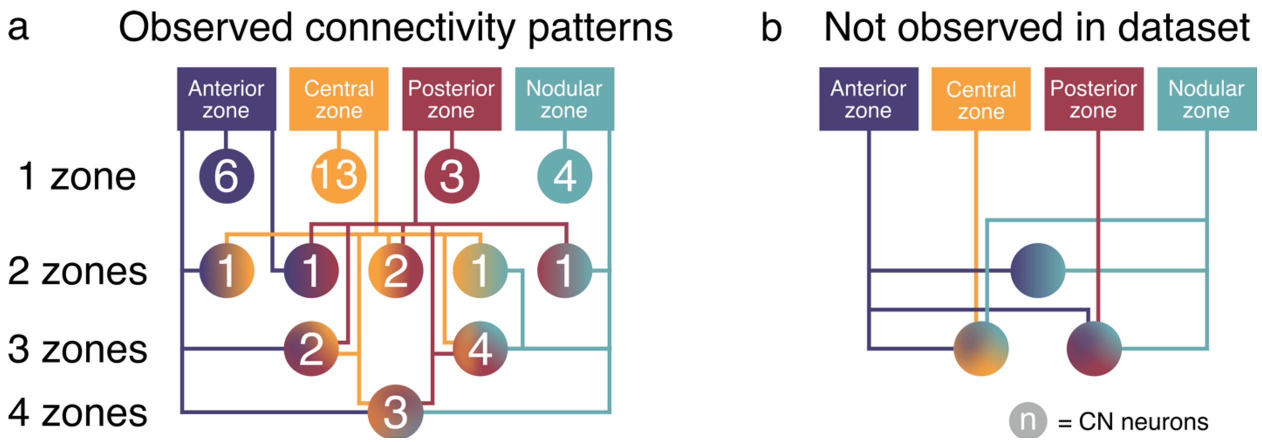


**Supplementary Figure 3**: **a**, Representation of observed Purkinje cell – CN neuron connectivity patterns based on cerebellar zones (circles). Numbers represent occurrences of each pattern in our dataset. **b**, The following connectivity patterns were not observed in our dataset: "AN," "ACN," and "APN" CN neurons.

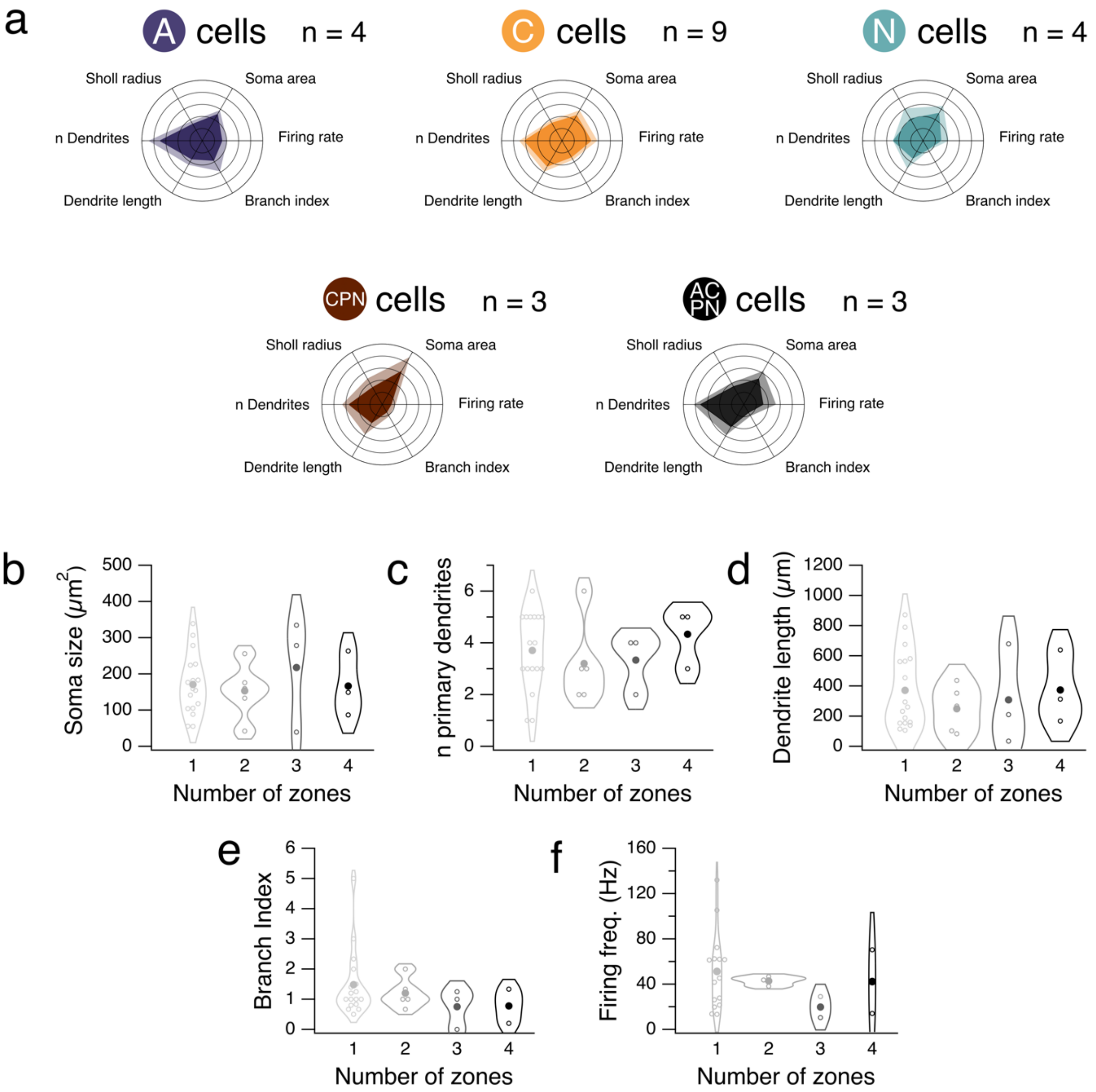


**Supplementary Figure 4**: Morphological and physiological properties of CN neurons. **a**, Polar plots displaying normalized average value of each measure per connectivity pattern for patterns with ≥ 3 cells per group. Shaded region indicates standard error. **b**, Soma size. **c**, Number of primary dendrites. **d**, Total dendrite length. **e**, Branch index at 25 µm. **f**, Firing frequency for cells by number of input zones. Unlabeled comparisons = not significantly different.

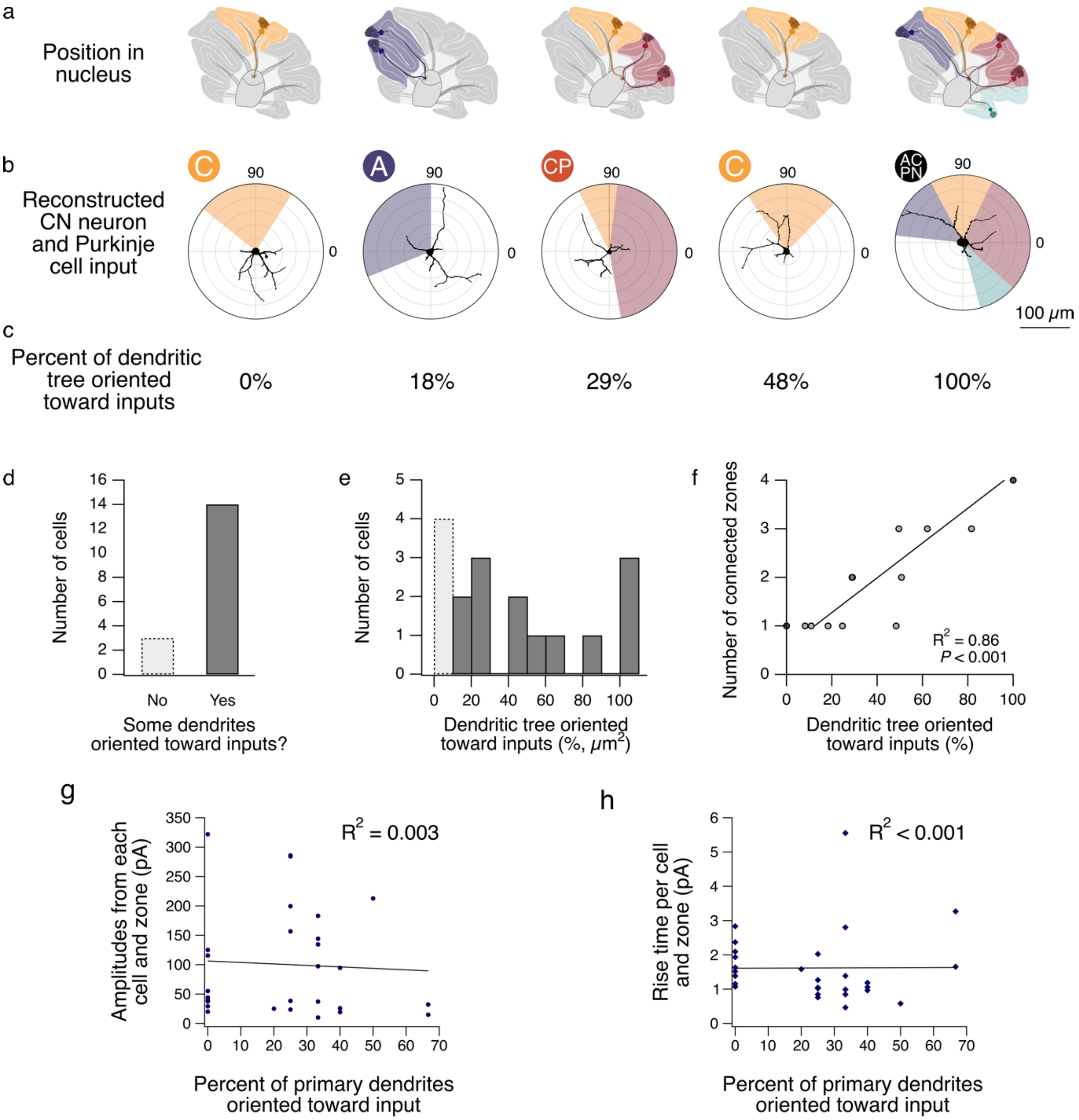


**Supplementary Figure 5**: CN dendrites are not preferentially oriented toward their zone inputs. **a**, Cartoons of cerebellar slices showing the relative position of individual CN neurons with different connectivity patterns. **b**, Polar plots showing the superimposed reconstructions of CN neurons from **a** with the input directions of the connected zones. **c**, Percent of dendritic tree for cells in **b** oriented toward their connected zones. **d,** Numbers of cells with and without dendrites oriented toward their inputs. **e**, Percent of CN neuron dendrite surface area oriented toward their connected zones, and **f**, as a function of the number of connected zones. Number of connected zones vs. percent of dendritic tree oriented toward inputs, Pearson's $r^2$ correlation coefficient. **g**, Amplitudes for each input per cell vs. the number of primary dendrites oriented toward that input zone (e.g. a four-zone cell has four data points). **h**, Same as **g**, but the rise time for each input.

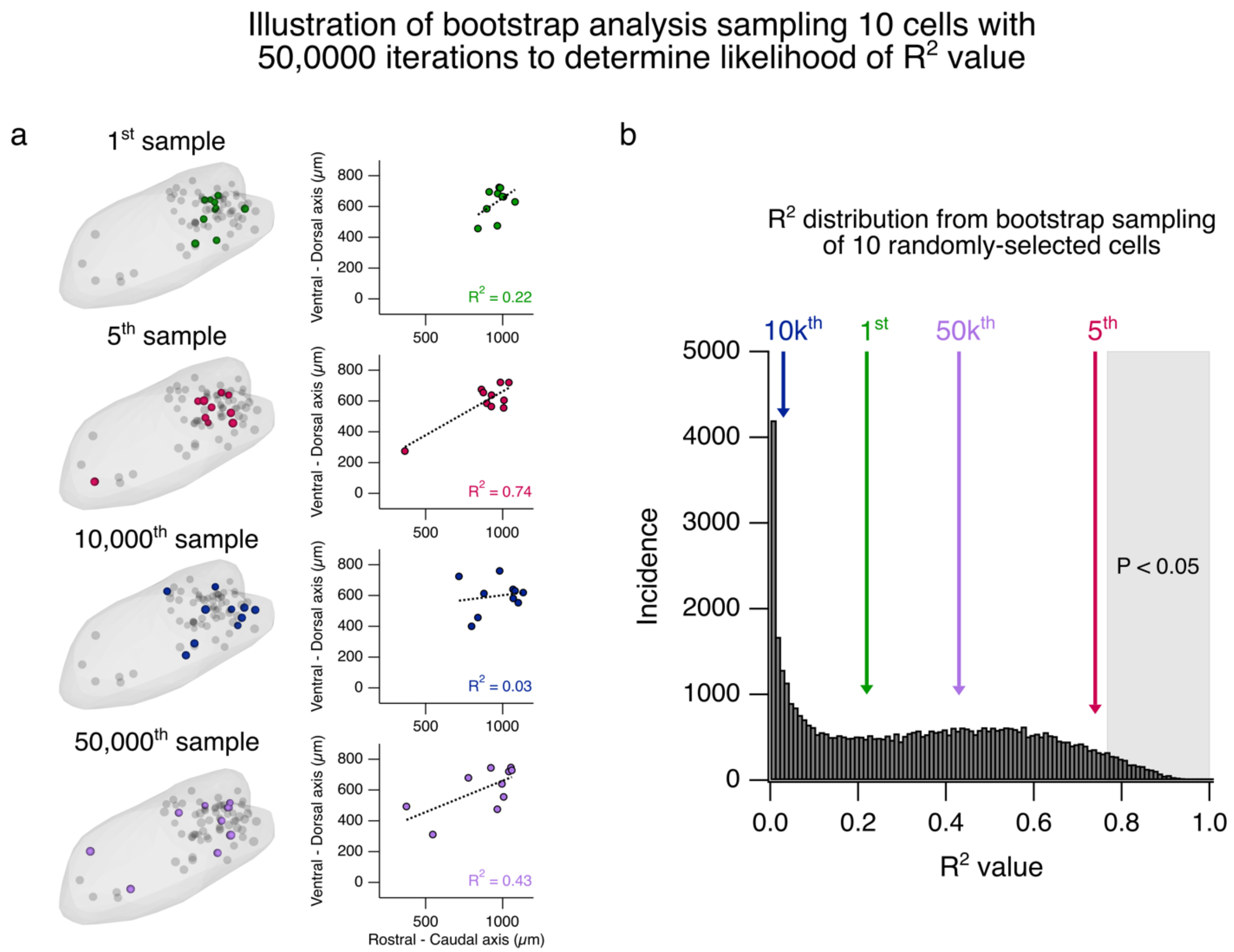


**Supplementary Figure 6**: Illustration of bootstrap analysis using 50,000 randomly selected samples of n cells to assess topographic organization within the CN. **a**, Left, renderings of CN in the sagittal plane depicting 10 randomly selected cells in the 1st, 5th, 10,000th, and 50,000th bootstrapped samples. Right, same as in **a, left,** but showing the plotted points for each CN neuron in the rostral-caudal and ventral-dorsal axes. Dotted lines and $R^2$ values in each graph represent the line of best fit for each sample of 10 cells. **b**, Distribution of $R^2$ values from the 50,000 bootstrapped samples. $R^2$ values from samples in **a** are highlighted with arrows. Shaded region depicts $R^2$ values that occurred in the top 5% of the distribution.

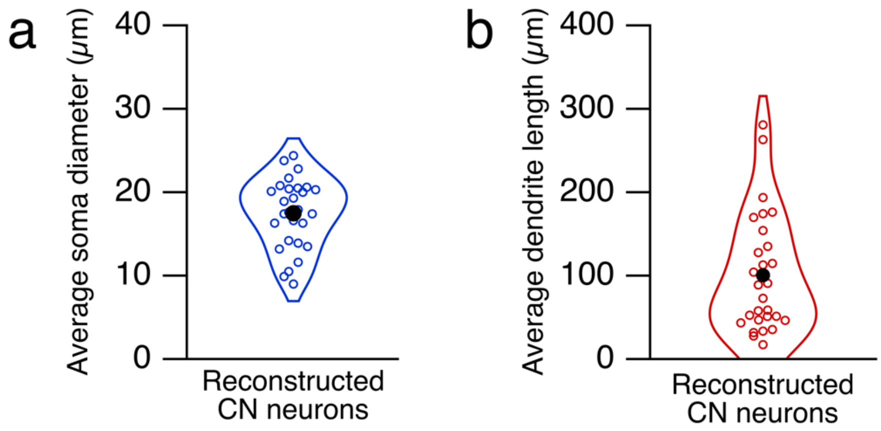


**Supplementary Figure 7**: Morphological properties from filled, reconstructed CN neurons. **a**, Average feret maximum soma diameter from Alexa-594-filled CN neurons. **b**, Average dendrite length from Alexa-594-filled CN neurons. n = 18 cells.

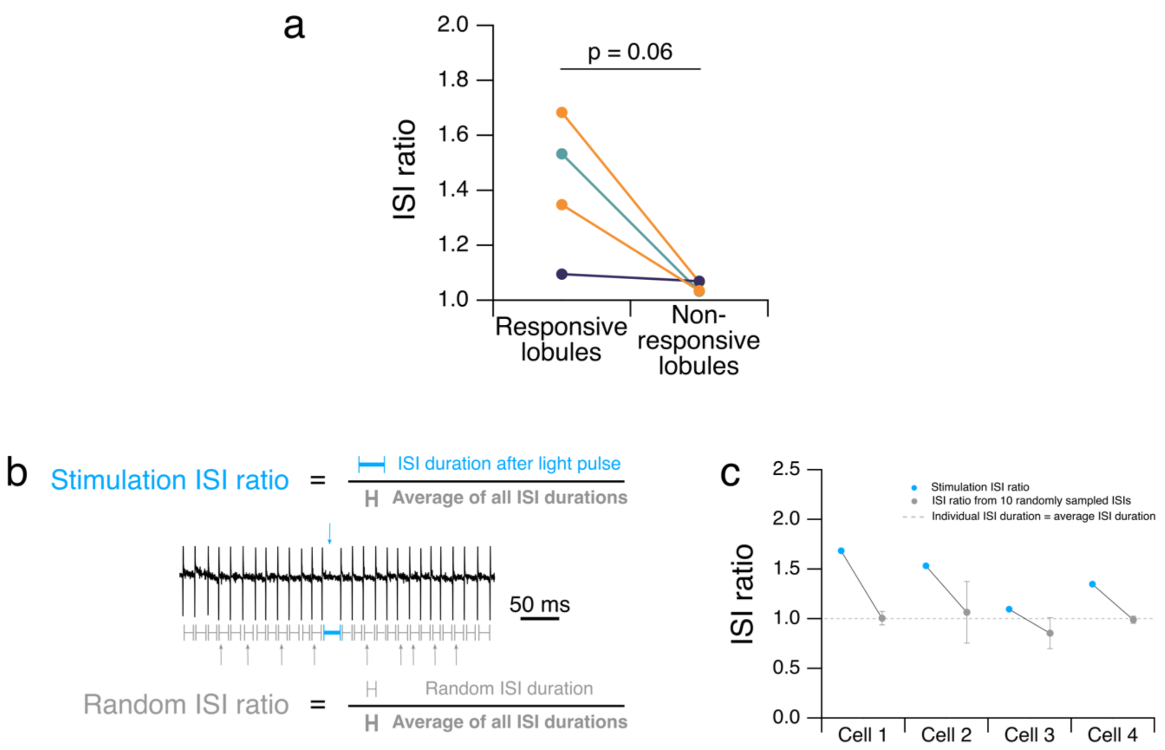


**Supplementary Figure 8**: The ISI ratio following stimulation of responsive lobules is higher in all tested cells compared to randomly sampled spike ISIs from the same trial. **a**, ISI ratios per cell for responsive lobules: 1.42 ± 0.25; non-responsive lobules: 1.05 ± 0.02. **b**, Top: ISI ratio calculated by dividing the ISI duration after stimulation with the average ISI duration per cell per trial. Middle: Example trace with blue and grey arrows highlighting the immediate post-stimulation ISI and other random ISIs, respectively. Bottom: Random ISI ratio calculated by dividing the ISI duration from a randomly sampled spike pause in the same trial as the stimulation, by the average ISI duration across the entire trial. **c**, Comparison of stimulation ISI ratios (blue) and 10 randomly selected ISI ratios (grey) for each cell. Error bars show the 95% confidence interval for ISI ratio acquired from the 10 randomly selected ISIs for each cell. Dotted line indicates ISI ratio of 1, where the selected ISI duration is equivalent to the average ISI duration for the entire trial. Unlabeled comparisons = not significantly different.

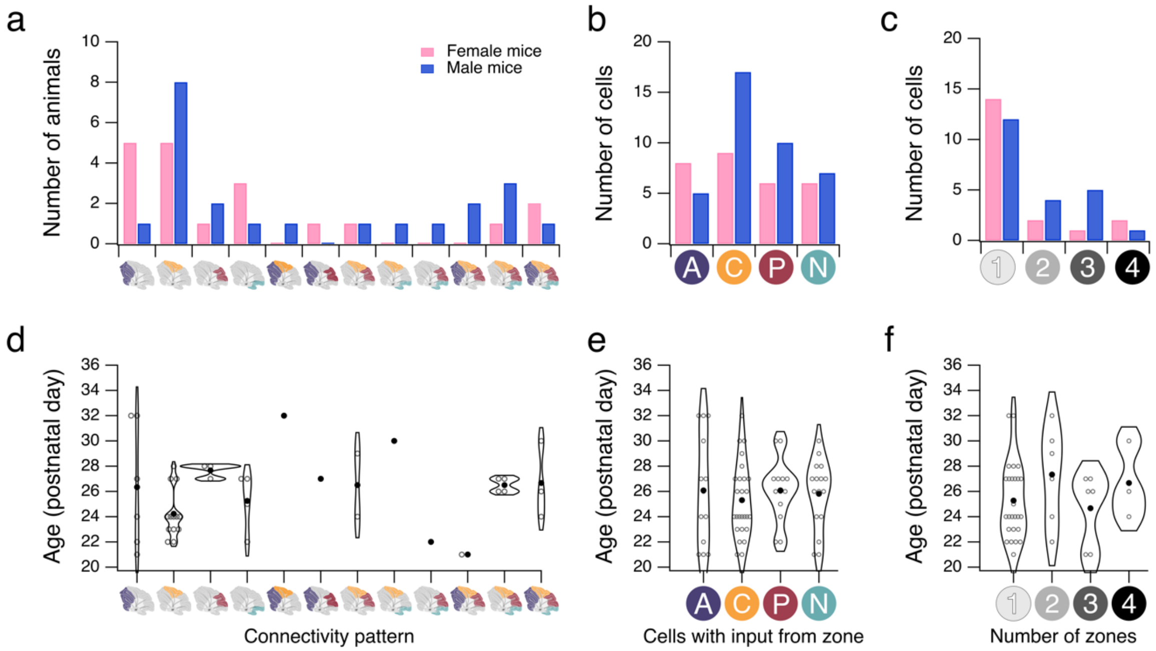


**Supplementary Figure 9**: No sex or age differences observed across different connectivity patterns. **a**, Distribution of observed connectivity patterns for female (pink) and male (blue) mice. **b**, Number of cells from female and male animals receiving input from each zone. **c**, Number of cells from female and male animals receiving input from n number of zones. **d**, Distribution of observed connectivity patterns across different ages. **e**, Distribution of different ages receiving input from each zone. **f**, Distribution of different ages receiving input from n number of zones. N = 22 female mice, 25 male mice. Unlabeled comparisons = not significantly different.

| Animal | Number of positive pixels in channel 1 | Number of positive pixels in channel 2 | Number of colocalized positive pixels | Percent colocalized pixels/total positive pixels |
|---|---|---|---|---|
| Animal_ID1 | 52351 | 6756 | 139 | 0.24 |
| Animal_ID2 | 75784 | 10140 | 88 | 0.10 |
| Animal_ID3 | 5758 | 4844 | 41 | 0.39 |
| Animal_ID4 | 107730 | 2808 | 157 | 0.14 |
| Animal_ID5 | 35239 | 12014 | 140 | 0.30 |
| Animal_ID6 | 8181 | 1392 | 0 | 0.0 |

**Table S1**: Colocalization of multiple viruses in virally labeled Purkinje cells. Number of positive pixels across imaging channels for six animals injected with fluorescently labeled viruses. Channels selected for colocalization analysis include the channel with the most viral expression (channel 1) and the channel with the second-most viral expression (channel 2). Numbers of positive pixels per channel represent the total number of positive pixels across >1 slices in imaging stack (2 μm interval).
